# Oxytocin modulation of self-other distinction is replicable and influenced by oxytocin receptor (OXTR) genotype

**DOI:** 10.1101/552703

**Authors:** Weihua Zhao, Ruixue Luo, Cornelia Sindermann, Jialin Li, Zhenyu Wei, Yingying Zhang, Congcong Liu, Jiao Le, Daniel S. Quintana, Christian Montag, Benjamin Becker, Keith M Kendrick

## Abstract

Intranasal oxytocin (OXT) has been associated with effects on diverse social-emotional domains in humans, however progress in the field is currently hampered by poor replicability. Limited statistical power and individual differences in biological factors, such as oxytocin receptor (OXTR) genetics, may have contributed to these variable findings. To this end, we present a pharmaco-genetic study aiming at (1) replicating previous findings suggesting that intranasal oxytocin (24 IU) blurs self-other distinction in a large sample of n = 170 male subjects, (2) determining whether variations in common receptor polymorphisms (rs237887, rs2268491, rs2254298, rs53576, rs2268498) influence sensitivity to oxytocin’s behavioral effects. Employing a validated oxytocin-sensitive trait judgment paradigm, we confirmed that it blurred self-other distinction in terms of decision time and subsequent memory. However, oxytocin only influenced decision time in rs53576 G carriers, whereas effects on memory performance were most pronounced in rs2268498 TT homozygotes. In summary, the current study replicates our previous findings showing that oxytocin blurs self-other distinction and suggests that sensitivity to its effects in this domain are receptor genotype dependent.

## INTRODUCTION

During the last decade the number of intranasal oxytocin (OXT) administration studies has steadily increased, with it being proposed to facilitate social interaction by increasing trust [1], emotional empathy [2], ‘mind-reading’ [3] and decreasing self-referential bias [4]. Such findings have inspired the hypothesis that OXT has the potential to alleviate social interaction deficits in psychiatric disorders and led to initial clinical trials in patients with some promising, but also variable therapeutic outcomes [5].

Despite initial promise, in recent years the field has been increasingly faced with critical evaluations of OXT’s effects on complex social behavior in humans (e.g. [6]; but see [7]). Particularly, low statistical power [8, 9] and replicability of findings [10] of intranasal OXT administration experiments have become a matter of growing concern. Indeed, recent comprehensive replication studies failed to replicate modulatory effects of intranasal OXT on core domains initially associated with OXT, including ‘mind reading’ [3, 11] as well as trust - both when money ([1]; for contrasting findings [12], critical review by [10]) or confidential personal information were at stake (see review [13]).

To explain the variable influence of OXT on social behavior, moderating effects of personal and contextual factors have increasingly moved into focus [8]. With respect to personal factors, converging evidence points to the important role of biological factors including sex [14, 15] and genetic variations, particularly individual differences in oxytocin-receptor gene (OXTR; located on chromosome 3p25) polymorphisms [16, 17].

Molecular genetic studies have reported associations between individual variations in single nucleotide polymorphisms (SNPs) of the OXTR gene and behavioral and neural differences in core social-emotional processes, including racial in-group bias (rs53576) [18], empathic reactivity (rs53576) [19], endorsement of prosocial behavior (rs2268498) [20]; as well as traits associated with social-emotional functioning, particularly autism-like traits (rs2268498) [21]. However, meta-analytic approaches have failed to confirm robust associations between the OXTR rs53576 and rs2254298 on social-emotional behavior [22], although a more recent meta-analysis demonstrated an association between OXTR rs53576 and empathy [23].

An increasing number of pharmaco-genetic studies has begun to explore whether individual differences in OXTR genetics influence the behavioral and neural sensitivity to intranasal OXT. Modulatory effects have been reported in the domains of facial emotion recognition [17, 24], social salience[16], and cooperation [14] suggesting that individual differences in OXTR genetics may account for the variable effects of intranasal OXT on interpersonal behavior.

Interpersonal behavior is shaped by perception of the interaction partner, yet also by the perceived differentiation between self and others [25]. The distinction between self and others is commonly operationalized as self-referential processing [26], that is biased processing of contents that are self-related, including biased evaluation and remembering personality traits pertaining to the self as opposed to others[4, 25]. Accumulating evidence suggests that OXT regulates self-referential processing [27, 28], which together with enhanced emotional empathy [2, 29] may underlie OXT facilitation of social interaction. For instance, in a previous study we combined intranasal administration of OXT with a self-referential task including self and other trait judgments and found that it reduced response times for both self and other trait judgments (i.e. facilitated decision making during judgments) but also decreased the accuracy of the subsequent recall of self-judgments, suggesting that it blurred the self-other distinction [4].

In the context of improving methodological standards in the field, particularly increased statistical power and replication, as well as increasing evidence for OXTR-modulated individual differences in the sensitivity to intranasal OXT effects, the present randomized placebo-controlled double-blind between-subjects pharmaco-genetic intranasal OXT administration study aimed to (1) determine the replicability of our previous findings (*n* = 38, male subjects) suggesting that intranasal OXT blurs the bias for self-referential trait judgments [4], in a cohort that is over four times larger (*n* = 170, male subjects), (2) determine whether the sensitivity to treatment effects in this domain is modulated by individual variations in OXTR genetics. We expected that (1) in line with our previous findings OXT would reduce reaction time for both self and other trait judgments and reduce their subsequent recall, (2) treatment effects would vary as a function of OXTR genotype.

## MATERIALS AND METHODS

### Participants

In a double-blind, between-subject placebo (PLC) controlled design, a total of 170 (*M*_age_ = 21.26 years, *SD* = 2.49 years) healthy male participants were randomly assigned to receive a single dose of 24 IU OXT (Sichuan Meike Pharmaceutical Co. Ltd, Sichuan, China; ingredients: oxytocin, glycerine, sodium chloride and purified water) (*n* = 86) or placebo (PLC, supplied by the same company with identical ingredients except for oxytocin) (*n* = 84). Treatment was administered following a standardized administartion protocol. The experimental asessments started 45 minutes after administration, the present paradigm was preceded by another behavioral task reported in a separate publication [30]. Participants could not determine better than chance whether they had received PLC or OXT (*χ*^2^ = 2.35, *p* = 0.13) confirming successful double blinding. Sample size was determined by estimating the power required to detect the primary outcome of interest, that is between-group differences (OXT vs. PLC) in reaction time and memory responses. Our previous study yielded between group differences in reaction times of *d* = 0.59 and memory of *d* = 0.95. Thus, we took a conservative approach and estimated a medium effect size to *d* = 0.5, which resulted in a requirement for 85 subjects to achieve 90% statistical power at *p* = 0.05. This level of statistical power (90%) is considerably higher than the average oxytocin study in healthy controls, which has been reported as 16% [9].

A total of *n* = 176 Chinese male (*M*_age_ = 21.33 years, *SD* = 2.48 years) were enrolled (Chengdu Gene Brain Behaviour Project, CGBBP). Participants were required to abstain from caffeine, alcohol, nicotine or other psychoactive substances in the 24h before the experiment. Exclusion criteria were any self-reported psychiatric illness, drug or alcohol abuse, left-handedness, or missing responses during the paradigm, leading to a sample size of *n* = 170 participants for the final analyses. The study was approved by the local ethics committee of the University of Electronic Science and Technology of China and the procedures were in accordance with the latest revision of the declaration of Helsinki. All subjects provided written informed consent and received monetary compensation for their participation.

### Experimental design, procedures and protocols

Before treatment, participants completed self-report questionnaires to account for potential between group differences in important confounders, including general confounders such as personality (NEO-Five Factor Inventory), depression(Beck Depression Inventory II), current mood and anxiety (Positive Affect and Negative Affect Schedule; State-Trait Anxiety Inventory), as well as traits with direct relevance to the paradigm (Inclusion of Others in Self; Self-Construal Scale). For genotyping all subjects additionally provided buccal swaps.

Subsequently, subjects were randomly allocated to receive either OXT or PLC and underwent a self vs. other trait judgment paradigm. Following a 4s instruction, subjects were asked to judge whether either a positive or negative adjective displayed described themselves, their mother (an extended self in Chinese culture) or a stranger. One of eight adjectives in each block (2 blocks in each judgment condition, i.e. 6 blocks in total) was presented (positive and negative words were balanced) and subjects made a judgment by button press (yes or no) within a time window of 2s. A ‘cue’ word (self, mother or a stranger) was displayed above the written trait adjective word (presented centered on the screen, **Figure 1**). In the subsequent “surprise” memory test, 24 trait adjectives (half positively and half negatively valenced) were randomly intermixed with 24 new trait adjectives, and subjects required to identify old vs. new items presented in a random order. The stimuli and design were identical to our previous study except for: (1) the number of blocks was decreased due to the high sensitivity observed in the previous study (2) additional conditions that served as baseline or control conditions during the previous fMRI study (classmate, font, and asterisk) were not included. All adjectives were displayed as two Chinese character words and balanced valence and arousal [4]. Based on our previous findings, primary outcome parameters to evaluate the effects of OXT were response time (RT) for the trait judgments and accuracy during the recognition memory test [4].

**Figure 1.**
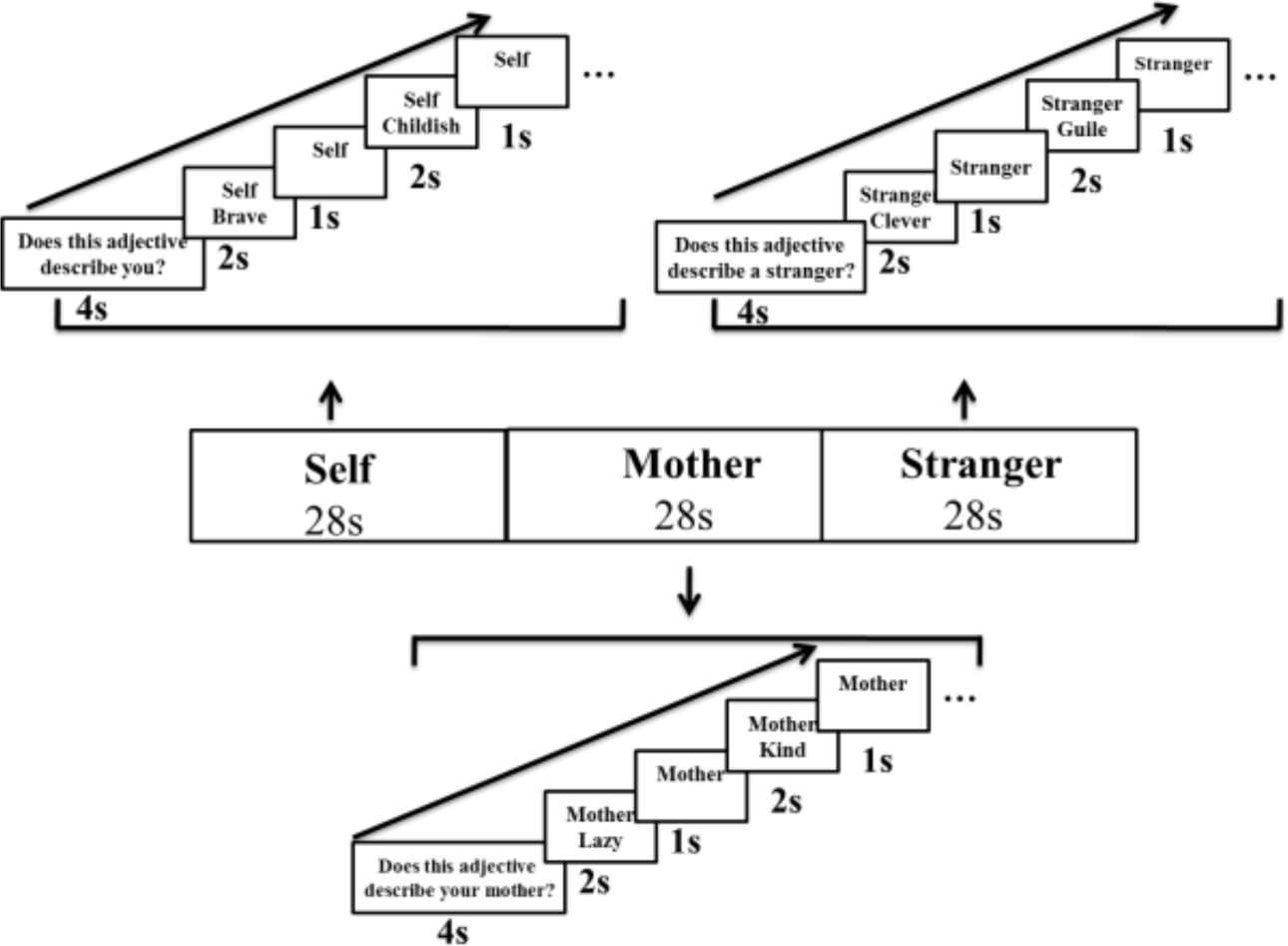
Trait-judgment task. A 4-s instruction that required subjects to judge if an adjective (either positive or negative valence) was appropriate to describe the person (either self, mother or a stranger) was displayed. Eight trait adjectives were presented in each session. Each trial consisted of a ‘cue’ word (either self, mother or a stranger) above a trait adjective presented for 2s at the center of the screen and it was followed by the cue for 1 s. The total time for each session was 28 s.

### Genotyping analyses

DNA was extracted from participants’ buccal cells. Automated purification of genomic DNA was conducted using a MagNA Pure 96 machine and commercial extraction kits (Roche Diagnostics, Mannheim, Germany). Five common OXTR SNPs were chosen based on previous studies suggesting associations with social behavior (OXTR rs237887, rs2268491, rs2254298, rs53576, rs2268498; e.g.[14]). Genotyping of the OXTR SNPs was performed by real-time polymerase chain reaction (PCR) and subsequent melting curve detection using a Cobas Z 480 Light Cycler (Roche Diagnostics, Mannheim, Germany). With the melting curve analyses, the alleles in each SNP were distinguished by different fluorescent labels of allele-specific oligonucleotide probe pairs. Simple probe assay designs from TIBMolBiol (Berlin, Germany) were used.

## RESULTS

There were no significant differences between the OXT (*n* = 86) and PLC (*n* = 84) treatment groups in terms of age, personality, mood, anxiety, depression or self-construct scores. Importantly, there were also no significant differences between scores on Inclusion of Others in Self for “mother” between the two groups (*p*s > 0.12) (**Table 1**).

**Table 1.** Age and questionnaire scores for study subjects in both groups (mean ± SD)

| Measurements | Placebo (*n* = 84) | Oxytocin (*n* = 86) | *t*-value | *p*-value |
| --- | --- | --- | --- | --- |
| Age (years) | 21.26±2.49 | 21.16±2.43 | 0.26 | 0.79 |
| Positive and Negative Affective Scale -Positive | 30.17±5.65 | 29.15±6.58 | 0.70 | 0.49 |
| Positive and Negative Affective Scale -Negative | 17.35±5.54 | 18.20±6.79 | 0.90 | 0.37 |
| Beck Depression Inventory | 5.90±6.91 | 6.12±7.34 | 0.19 | 0.85 |
| State-Trait Anxiety Inventory -State | 38.04±7.88 | 38.39±9.04 | 0.27 | 0.79 |
| State-Trait Anxiety Inventory -Trait | 41.00±7.76 | 40.50±8.03 | 0.41 | 0.69 |
| NEO-Five Factor Inventory |  |  |  |  |
| Neuroticism | 33.95±7.15 | 34.21±7.30 | 0.23 | 0.82 |
| Extraversion | 38.38±5.22 | 38.86±5.67 | 0.57 | 0.57 |
| Openness to experience | 39.80±5.48 | 39.58±5.13 | 0.27 | 0.79 |
| Agreeableness | 42.14±4.96 | 41.81±5.30 | 0.42 | 0.68 |
| Conscientiousness | 40.68±5.01 | 40.96±5.55 | 0.35 | 0.73 |
| Self-Construal Scale -independent | 72.96±9.69 | 72.81±10.05 | 0.09 | 0.93 |
| Self-Construal Scale -interdependent | 74.42±9.34 | 75.63±9.92 | 0.73 | 0.47 |
| Inclusion of Others in Self -mother | 4.87±1.29 | 4.57±1.19 | 1.57 | 0.12 |

### Distribution of genotypes

In the current sample, distribution of SNP genotypes was in the Hardy–Weinberg equilibrium (HWE); for rs237887, GG = 55, GA = 85, AA = 30 [*n* = 170, HWE: *χ*^2^ = 0.08, *p* =0.77]; for rs2254298, GG = 79, GA = 75, AA = 16 (*n* = 170, HWE: *χ*^2^ = 0.09, *p* = 0.77); for rs2268491, CC = 80, CT = 74, TT = 15 (*n* = 169, HWE: *χ*^2^ = 0.13*, p* = 0.72); for rs2268498, CC = 11, CT = 74, TT = 84 (*n* = 169, HWE: *χ*^2^ = 0.99, *p* = 0.32); for rs53576, AA = 87, AG = 65, GG = 13 (*n* = 165, HWE: *χ*^2^ = 0.03, *p* = 0.86).

To increase statistical power and avoid statistical inference errors due to the small sample size of the GG group, we grouped the alleles into G+ (GG/AG, 78 males: 40 in OXT group, 38 in PLC group) and G- (AA, 87 males: 43 in OXT group, 44 in PLC group) for rs53576 in the judgment task analysis [31]. In addition, there were no significant differences between G+ and G- allele groups across OXT and PLC groups in all questionnaires mentioned above (*p*s > 0.09), indicating confounders were controlled for. For rs2268498, we grouped the alleles into C+ (CC/ CT, 83 males: 40 in OXT group, 43 in PLC group) and C- (TT, 83 males: 44 in OXT group, 39 in PLC group) in the memory performance task analysis. No significant group differences between C+ and C- allele groups across OXT and PLC groups were found in confounding factors (*p*s > 0.11).

### Intranasal OXT effects on self-other distinction

In an initial analysis step the replicability of the effects of intranasal OXT on self- other processing independent of genotype was evaluated. We performed a mixed three-way ANOVA with treatment (OXT vs. PLC) as between-subject factor, condition (self, mother, stranger) and valence (positive vs. negative) as within-subject factors, and RT as dependent variable. Results revealed a significant main effect of condition [*F*_(2,336)_ = 21.67, *p* < 0.001, *η*^2^ = 0.11] suggesting faster self (*p* < 0.001, Cohen’s *d* = 0.48) and mother (*p* < 0.001, Cohen’s *d* = 0.36) judgments compared with stranger judgments across the two treatment groups, as well as significant main effects of valence [*F*_(1,168)_ = 16.12, *p* < 0.001, *η*^2^ = 0.09] and treatment [*F*_(1,168)_ = 5.55, *p* = 0.02, *η*^2^ = 0.03]. In addition, condition x treatment [*F*_(2,336)_ = 6.46, *p* = 0.002, *η*^2^ = 0.04] and condition x valence [*F*_(2,336)_ = 18.25, *p* < 0.001, *η*^2^ = 0.10] interactions were significant. Post-hoc analysis of the interaction effect between condition and treatment with Bonferroni correction suggested that OXT decreased RT across all conditions but only reached significance in the stranger condition (*p* = 0.001, Cohen’s *d* = 0.54). In the PLC group, RTs for judging stranger’s traits were slower than for self (*p* < 0.001, Cohen’s *d* = 0.65) and for mother (*p* < 0.001, Cohen’s *d* = 0.62, **Figure 2**). In the OXT group there was only a significant difference between self and stranger (*p* = 0.013, Cohen’s *d* = 0.30).

**Figure 2.**
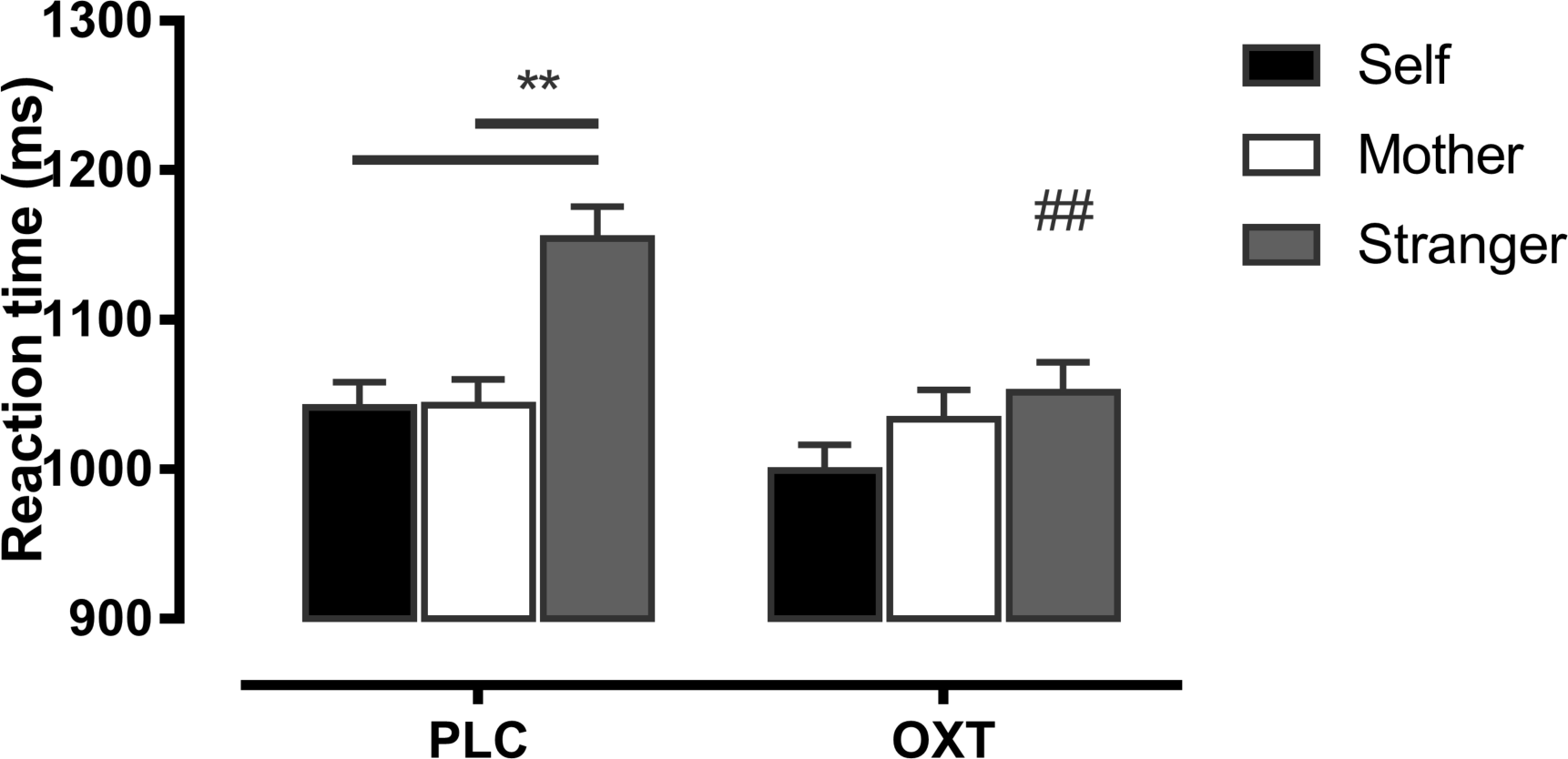
Histograms show mean ± sem reaction time for judging self-, mother-, stranger-related adjectives following OXT and PLC administration. \*\**p* < 0.01 for differences within each group; ^##^*p* < 0.01 for OXT vs. PLC.

Due to technical problems (missing responses), three subjects (*n* = 1 OXT, *n* = 2 PLC) were additionally excluded from the subsequent memory analysis. A concordant three-way mixed ANOVA with recognition accuracy as dependent variable revealed significant main effects of condition [*F*_(2,330)_ = 74.14, *p* < 0.001, *η*^2^ = 0.31, reflecting better memory performance in self condition compared to mother condition (*p* < 0.001, Cohen’s *d* = 0.77) and stranger condition (*p* < 0.001, Cohen’s *d* = 1.07)] and valence [*F*_(1,165)_ = 14.76, *p* < 0.001, *η*^2^ = 0.08, due to higher accuracy to positive vs. negative valence adjectives (*p* < 0.001, Cohen’s *d* = 0.38)]. In addition, there was a significant two-way interaction between condition and valence [*F*_(2,330)_ = 5.87, *p* = 0.003, *η*^2^ = 0.03]. Based on the findings from our previous study demonstrating an effect of OXT on self-referential memory we specifically explored this effect in the treatment groups separately. The results suggested higher accuracy for recognizing both, positively and negatively valenced adjectives for self as compared to mother (*ps* < 0.001) or stranger (*ps* < 0.001) conditions. In contrast there was no difference between mother and stranger conditions for recognition accuracy of negative trait adjectives following OXT (*p* = 0.62).

To further examine whether the observed effects of OXT on RT and accuracy were associated, a Pearson correlation was conducted but no significant associations occurred within the treatment groups (*p*s > 0.10), suggesting that effects on the two domains of self- versus other processing were independent. However, in contrast with our previous findings no significant three-way interaction between condition, valence and treatment was observed [*F*_(2,330)_ = 0.79, *p* = 0.45].

### Interaction effects between OXTR genetics and intranasal OXT effects

To identify the modulation of OXTR genetic variations, OXTR genotype was included as an additional between-subject factor in the subsequent mixed ANOVA models.

For RT, we did not find any significant interaction effects across treatment, genotype, condition and valence for OXTR rs2254298, rs2268491, rs237887, rs2268498 (*F*s <1.06, *ps* > 0.38), however a significant four-way interaction effect was found for rs53576 [*F*_(4,318)_ = 3.91, *p* = 0.004- (*p* _Bonferroni (corrected for SNPs_ = 0.02)]. For AA genotype subjects of this SNP significant main effects of condition [*F* _(2,170)_ = 10.79, *p* < 0.001, *η*^2^ = 0.11] and valence [*F*_(1,85)_ = 12.54, *p* < 0.001, *η*^2^ = 0.13] and an interaction between the two factors [*F*_(2,170)_ = 14.77, *p* < 0.001, *η*^2^ = 0.15] were observed. On the other hand, for the G+ allele subgroup of OXTR rs53576, a main effect of condition [*F*_(2,152)_ = 12.73, *p* < 0.001, *η*^2^ = 0.14], valence [*F*_(1,76)_ = 4.93, *p* = 0.03, *η*^2^ = 0.06], and a two-way interaction effect between condition and valence [*F*_(2,152)_ = 9.03, *p* < 0.001, *η*^2^ = 0.11] reached significance. Notably, we also found a two-way interaction effect between treatment and condition [*F*_(2,152)_ = 4.15, *p* = 0.018, *η*^2^ = 0.05] and a three- way interaction across treatment, condition and valence [*F*_(2,152)_ = 5.35, *p* = 0.006, *η*^2^ = 0.07]. Bonferroni-corrected post-hoc tests of the two-way interaction effect suggested that in the PLC group exhibited slower RT judging trait adjectives of stranger compared to self (*p* < 0.001, Cohen’s *d* = 0.74) or mother (*p* < 0.001, Cohen’s *d* = 0.64), however we did not find any other significant effects in the OXT group (all *p* > 0.29). A significant difference in RT between OXT and PLC in the stranger condition was also found [*p* = 0.002, Cohen’s *d* = 0.73] (**Figure 3A**).

**Figure 3.**
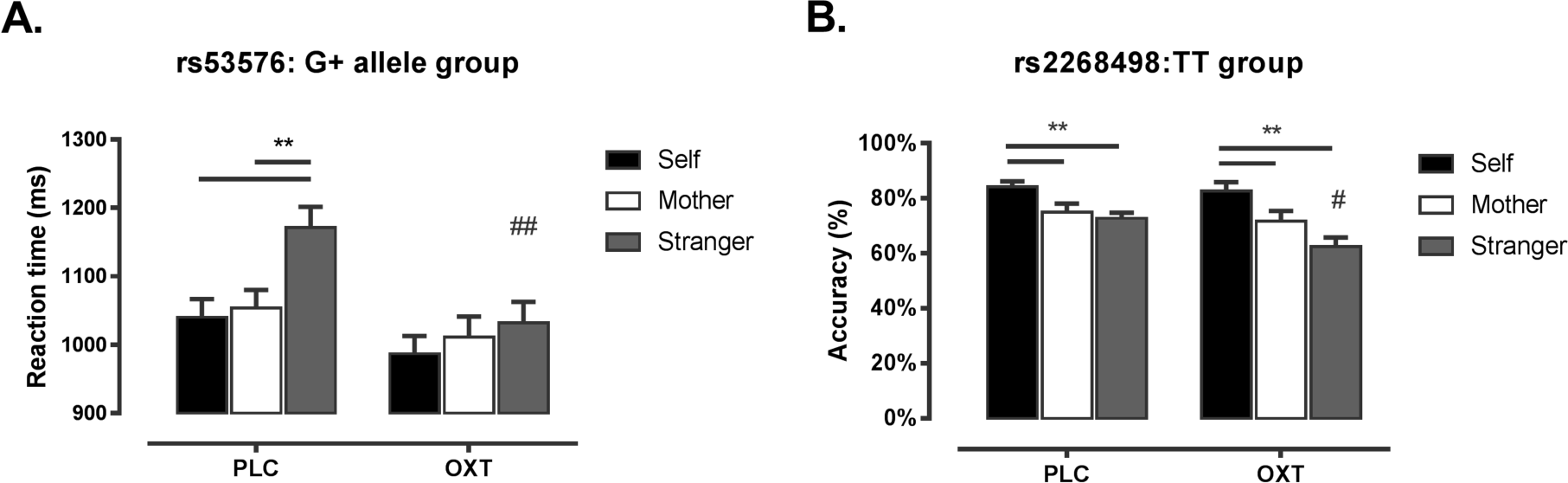
(A) Histograms show mean ± sem reaction time for judging self-, mother-, stranger-related adjectives in OXT and PLC subgroups for individuals with G+ allele of OXTR rs53576 (B) Histograms show mean ± sem memory performance for remembering self-, mother-, stranger- related negative words in OXT and PLC for individuals with TT genotype of OXTR rs2268498. \*\**p* < 0.01 for differences within each group; ^#^*p* < 0.05 for OXT vs. PLC; ^##^*p* < 0.01 for OXT vs. PLC.

A four-way mixed ANOVA examining subsequent memory accuracy revealed no significant interaction effects across treatment, genotype, condition and valence for OXTR rs2254298, rs2268491, rs237887, rs53576 (*F*s < 1.60, *p*s > 0.17), although a significant four-way interaction effect was found for rs2268498 [*F*_(2,324)_ = 4.66, *p* = 0.01- (*p* _Bonferroni_ = 0.05)]. For rs2268498 C+ carriers significant main effects of condition [*F*_(2,162)_ = 33.73, *p* < 0.001, *η*^2^ = 0.29] and valence [*F*_(1,81)_ = 7.78, *p* = 0.007, *η*^2^ = 0.09] were observed. For the TT genotype, main effects of condition [*F*_(2,162)_ = 43.28, *p* < 0.001, *η*^2^ = 0.35] and valence [*F*_(1,81)_ = 6.99, *p* = 0.01, *η*^2^ = 0.08] as well as two- way interaction effect between condition and valence [*F*_(2,162)_ = 3.75, *p* = 0.03, *η*^2^ = 0.04] reached significance. Contrary to the non-significant three-way interaction effect in the entire sample, we found a significant three-way interaction effect across condition, valence and treatment [*F*_(2,162)_ = 3.10, *p* = 0.05, *η*^2^ = 0.04]. Bonferroni corrected post- hoc analysis suggested that OXT decreased memory accuracy for negatively valenced trait adjectives for ‘stranger’ relative to PLC [*F*_(1,81)_ = 6.31, *p* = 0.01, Cohen’s d = 0.55] (Figure 3B). In addition, accuracy of remembering trait adjective for both valences in the self condition was better than for mother and stranger conditions in PLC and OXT groups (all *p*s < 0.02). Correlation analysis between RT for judging words and accuracy of subsequent memory in these two different OXTR genotype groups across OXT and PLC (*p*s > 0.08) again did not reveal significant associations. Together, the results suggest that the effects of intranasal OXT on RTs may have been driven by pronounced effects in G carriers of OXTR rs53576, whereas effects on memory performance may have been driven by TT carriers of OXTR rs2268498.

## DISCUSSION

The present study aimed at determining the robustness of effects of intranasal OXT on self-other processing as well as their modulation by individual differences in OXTR genotype. Combining a previously validated and intranasal OXT-sensitive self-referential paradigm with a behavioral genetic approach in a large sample (*n* = 170) with sufficient statistical power allowed us to replicate that intranasal OXT abolished the self-referential bias in terms of reaction times during trait judgments [4]. Sensitivity to the behavioral effects of OXT on the self-referential bias was also modulated by individual differences in OXTR genotype. Trait judgment RTs were reduced more in G carriers of rs53576 relative to the AA genotype, whereas subjects homozygous for the T allele of rs2268498 showed pronounced effects on subsequent memory performance compared to C carriers.

In line with our previous findings [4], intranasal OXT blurred the self-other distinction by decreasing RTs for judging traits of a stranger. Although initial studies reported that intranasal OXT may enhance positive self-evaluation [32] or increase differential neural processing of self vs. celebrity judgements [33], the present findings demonstrated that OXT may rather attenuate the self-referential processing bias. These findings are in line with a growing number of recent studies reporting that intranasal OXT promotes self-other integration [34], facilitates perception of other- but not self-experienced pain [35], and increases other-orientation [36]. Studies that concurrently assessed eye gaze or neural activity suggest that OXT-enhanced other orientation specifically affects implicit processing [37] and is mediated by modulatory effects on brain regions engaged in self-referential processes, particularly the medial prefrontal cortex [4, 38].

An overarching aim of the present study was to determine the robustness of OXT’s effects on self-referential processing using a sufficiently powered design [8–10]. Despite failed replication attempts of OXT’s modulatory effects in the domains of ‘mind reading’ [3, 11] and trust [1, 10] the present findings resonate with recent findings demonstrating the robustness of OXT-enhanced emotional empathy [2]. It is unlikely that OXT exerts effects on all aspects on social cognition, thus it is important to identify which specific aspects it does modulate robustly. Together with the present results, these findings indicate an important role of OXT in increasing other-orientation which may represent a common denominator underlying its potential to facilitate social interaction.

In line with increasing evidence for the modulation of intranasal OXT effects by biological factors including OXTR genetics [16, 17] and that OXTR expression patterns in the human brain are associated with social cognition [39], a second aim of the present study was to determine whether the sensitivity to the behavioral effects of intranasal OXT varies as a function of individual differences in OXTR genotype. Of the five common OXTR variants that were examined only rs53576 and rs2268498 modulated individual sensitivity to intranasal OXT. Specifically, in G carriers of rs53576 accelerated trait judgements were observed in the non-self referential condition, suggesting that subjects with this polymorphism may have an increased sensitivity to intranasal OXT effects on speed of making social evaluation decisions. The OXTR rs53576 SNP has been associated with affiliative behaviour [40], with GG carriers demonstrating more sensitive parental interaction [41], continuous attachment security [42], higher sociability [43] and empathy [19] as well as higher psychological resources and prosocial temperament [44]. Together with a previous report on pronounced effects of intranasal OXT-enhanced preference for highly salient social stimuli (infant faces) [45] and social cooperation in male participants [14] as well as neural and behavioural effects in autistic individuals [46], the present findings suggest that the G allele promotes effects of intranasal OXT on other orientation. While G- carriers have primarily been associated with prosocial attributes, A- carriers have been associated more with impaired social attributes in the context of autism and empathy [47]. A study reported that intranasal oxytocin suppression of amygdala responses to fear faces and behavioural ratings of fear intensity to were restricted to AA homozygotes of OXTR rs53576 [24]. This may reflect a greater association of OXT effects on the amygdala in A-carriers and those on the medial prefrontal cortex on G-carriers. Thus, specific OXTR genotype modulation of intranasal OXT effects may be both region and task specific.

Although previous findings on interaction effects involving self-other condition, valence and treatment during the subsequent recognition memory test in the paradigm could not be replicated [4], including OXTR genotype as an additional factor in the analysis revealed that intranasal OXT decreased memory performance for negatively valenced words during the stranger condition specifically for rs2268498 T homozygotes but not C carriers. Previous studies reported that the rs2268498 T homozygotes had higher scores of empathic concern relative to those carrying a C allele [20] and that T-allele variant associates with better social perception [48] and lower trait autism scores [21]. On the other hand, C carriers have a greater association with higher trait autism [21].

Interestingly, OXT effects on trait decisions and subsequent memory were found to be modulated by different OXTR polymorphisms. In accordance with the finding that effects of OXT on the two domains of self-referential processing were not associated [4] this may suggest that differential mechanisms of OXT and associated OXTR mediated sensitivity may underlie the observed pattern of effects. Indeed, previous studies suggest that both types of processing rely on different neural systems, with effects of intranasal OXT on self-referential processing being mediated by modulatory effects on the medial prefrontal cortex [4, 38], whereas OXT effects on emotional modulation of memory formation may be mediated by the insula [49].

Some limitations in the current study need to be considered. Although the results demonstrate robust effects of exogenous OXT on self-referential processing in Chinese men, the replication was carried out by the same research group and within highly similar participant samples. To further increase trust in replicable and general effects of intranasal OXT future studies should aim at independent replications across laboratories, sexes and cultures (initial attempts to generalize treatment effects across cultures in [2]; or cross-cultural replication of OXTR associations with social behaviour in [21]).

Overall, the present study replicated previous findings on the effects of intranasal OXT on self- and other- processing, suggesting that OXT blurs self and other distinction. Individual differences in OXTR genotype determined sensitivity to the behavioural effects of OXT. The present findings add to the growing literature on an important role of the OXTR in modulating the effects of OXT administration [24, 50] and thus may help to identify individuals with a high responsiveness to OXT treatment in disorders characterized by altered self-referential processing.

## Funding and disclosure

This work was supported by grants from National Natural Science Foundation of China (NSFC) [31530032; 91632117]; Fundamental Research Funds for the Central Universities [ZYGX2015Z002]; Science, Innovation and Technology Department of the Sichuan Province [2018JY0001]; Open Research Fund of the State Key Laboratory of Cognitive Neuroscience and Learning (Beijing Normal University); The Novo Nordisk Foundation (NFF160C0019856).

The authors declare no competing financial interests.

